# GDF15 is a Critical Renostat in the Defense Against Hypoglycemia

**DOI:** 10.1101/2023.06.22.546095

**Authors:** Zongyu Li, Xinyi Zhang, Wanling Zhu, Qian Xu, Chuyue Yu, Qiancheng Zhao, Stephan Siebel, Cuiling Zhang, Bruna Genisa Costa Lima, Xiruo Li, Kavita Israni-Winger, Ali R. Nasiri, Michael J. Caplan, Rui B. Chang, Andrew Wang, Renata Belfort de Aguiar, Janice J. Hwang, Rachel J. Perry

## Abstract

Episodic hypoglycemia is one of the best honed, evolutionary conserved phenomena in biology, because of the constant feast-fast cycles that have characterized most of history. The counterregulatory response to hypoglycemia, mobilizing substrate stores to produce glucose, is the primary adaptive mechanism to enable survival. Catecholamines and glucagon have long been considered the key hypoglycemia counterregulatory hormones, but here we identify a new hypoglycemia counterregulatory factor. We employed the insulin tolerance test (ITT) and hyperinsulinemic-hypoglycemic clamp to mimic the two common settings in which hypoglycemia can occur in patients: postprandial insulin overdose and elevated basal insulin administration, respectively. We found that Growth Differentiation Factor 15 (GDF15) production is induced in the S3 segment of the renal proximal tubules and its release increases hepatic gluconeogenesis by increasing intrahepatic lipolysis in a beta-adrenergic receptor-2 (Adrb2)-dependent manner. In addition, mice exposed to recurrent hypoglycemia and patients with T1D exhibit impaired GDF15 production in the setting of hypoglycemia. These data demonstrate that GDF15 acts acutely as a gluco-counterregulatory hormone and identify a critical role for kidney-derived GDF15 in glucose homeostasis under physiological and pathophysiological conditions.

## Introduction

The defense against severe hypoglycemia under fasting conditions is of indispensable evolutionary importance. Therefore, it is logical that the mechanisms of hypoglycemia counterregulation are redundant and require a coordinated response of several hormones, including corticosteroids, catecholamines, glucagon, growth hormone – and perhaps others.

While insulin-treated diabetes has only existed for a century, and thus does not have the evolutionary gravity of fasting, in recent years the occurrence of insulin-induced hypoglycemia has gained clinical import. Since the 1970s, consistently improving glucose-monitoring technology has made tight glycemic control possible, even in those with type 1 and advanced type 2 diabetes requiring exogenous insulin. Aggressive insulin treatment is beneficial in reducing the risk of lifespan- and healthspan-limiting diabetes complications, including neuropathy, nephropathy, and cardiovascular disease ^1^. However, strict diabetes control is not without risks: in patients with T1D, therapeutic insulin may cause hypoglycemia: 35-42% of type 1 diabetes (T1D) patients suffer from severe hypoglycemia ^2^ and the risk of hypoglycemia increases with the duration of diabetes ^3^. The Action in Diabetes and Vascular Disease: Preterax and Diamicron Modified Release Controlled Evaluation (ADVANCE) trial demonstrated a 2-3-fold increased risk of major vascular events and of cardiovascular events following a single episode of severe hypoglycemia in patients pursuing aggressive glucose control for their diabetes ^4^.

Except for the brain, the influx and efflux of glucose in tissues are precisely regulated by hormones including insulin, glucagon, cortisol, epinephrine, and growth hormone. Conventional wisdom holds that hypoglycemia first triggers the cessation of β-cell insulin secretion, followed by secretion of glucagon, epinephrine, and cortisol. However, we determine in the current study that this list is not exhaustive: here we identify GDF15 as a novel hypoglycemia counterregulatory factor. GDF15, a member of the transforming growth factor-beta (TGFb) superfamily, is induced by various forms of cellular stress, including mitochondrial stress ^5^, liver injury ^6^, and ER stress ^7^, as well as with smoking, age, and exercise ^8^, Acutely, GDF15 can be induced both by bacterial and viral inflammation^9^ and by treatment with a subset of non-steroidal anti-inflammatory agents ^10^. In this study, we found that GDF15 is also induced in the setting of both insulin- and fasting-induced hypoglycemia.

To address the role of GDF15 in the setting of hypoglycemia, we applied state-of-the-art stable isotope tracer methodology in multiple strains of knockout and reporter mice. We reveal here that GDF15 increases rates of gluconeogenesis by activating intrahepatic lipolysis in response to Adrb2. However, the counterregulatory effect of GDF15 is attenuated in mice and humans with type 1 diabetes. Together, these data identify GDF15 as a new hypoglycemia counterregulatory hormone and provide insight into additional mechanisms of regulation of endogenous glucose production under physiologic and pathophysiologic conditions.

## Results

### Hypoglycemia – and impaired intracellular glucose metabolism – can induce GDF15 production

To generate hypoglycemia under clinically relevant conditions, we utilized two rodent models: the hyperinsulinemic-hypoglycemic clamp, mimicking increased basal insulin infusion; and the insulin tolerance test (ITT), mimicking excess prandial insulin dosing. As compared to rats undergoing a hyperinsulinemic-euglycemic clamp, hypoglycemic rats exhibited a 10-fold increase in plasma GDF15 concentrations (Fig. 1A-B). To validate this result in a second species, we performed hyperinsulinemic-hypoglycemic clamps in mice and found a similar induction of GDF15 by hypoglycemia (Fig. 1C-D). Next, to determine whether hypoglycemia from a single insulin overdose can induce GDF15 production, we performed an ITT in mice. Similar to the hypoglycemic clamp, ITT also induced an increase in plasma GDF15, but with a relatively lower amplitude (Fig. 1E-F). Compared to the male mice, age-matched female mice showed a lower basal GDF15 (Supplemental Fig. 1A-B) and similar peak GDF15 during ITT (Supplemental Fig. 1C), resulting in a greater degree of GDF15 induction (Supplemental Fig. 1D). However, to avoid any confounding from estrus cycling, we studied males in all of the subsequent preclinical studies in this manuscript, reasoning that because if anything GDF15 induction was lower in males than females, studies focusing on males would not overstate, but if anything would understate, the physiological role of GDF15 in hypoglycemia counterregulation.

**Figure 1.**
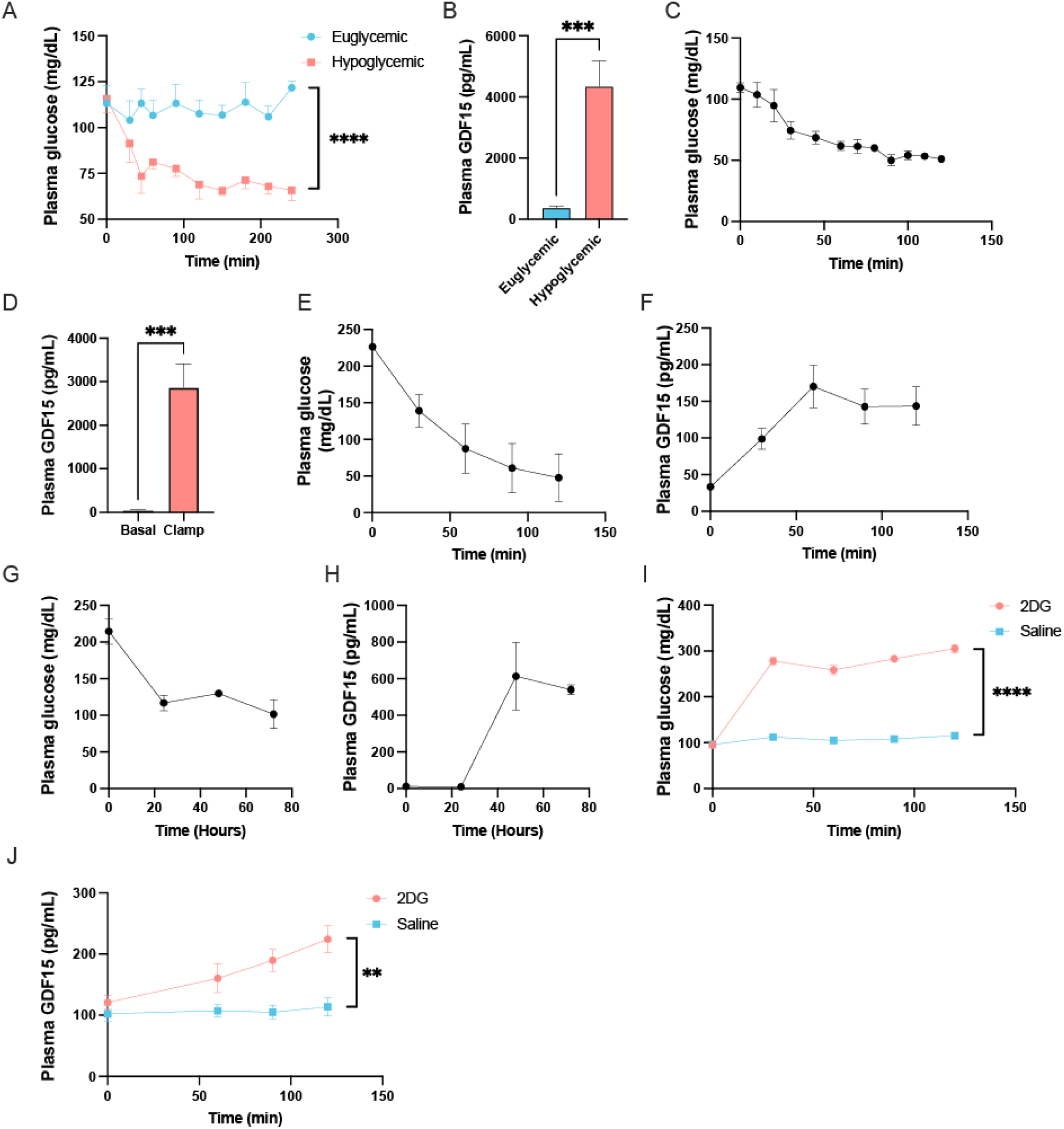
Hypoglycemia induces GDF15 production in mice and rats. (A) Plasma glucose and (B) GDF15 concentrations in rats undergoing hyperinsulinemic-euglycemic or - hypoglycemic clamps. (C) Plasma glucose and (D) GDF15 concentrations in mice during a hypoglycemic clamp. (E) Plasma glucose and (F) GDF15 concentrations in mice during an insulin tolerance test in mice. (G) Plasma glucose and (H) GDF15 concentrations in mice during a 72 hour fast in mice. (I) Plasma glucose and (J) GDF15 concentrations after treatment with 2-deoxyglucose (2DG). In all panels, \*\**P*<0.01, \*\*\**P*<0.001, \*\*\*\**P*<0.0001 by the 2-tailed unpaired Student’s t-test.

To interrogate if GDF15 production can be induced by insulin-independent hypoglycemia, we fasted healthy mice for 72 hours. Consistent with a previous report, we found serum GDF15 remained unchanged after 24 hours of fasting ^11^; however, GDF15 was robustly induced after 48 hours (Fig. 1H). To determine whether the induction of GDF15 during hypoglycemia required reductions in circulating glucose *per se*, we treated mice with 2-deoxy-D-glucose (2DG) to inhibit intracellular glucose metabolism without causing systemic hypoglycemia. After 2DG treatment, blood glucose concentrations increased within 30 min, indicating a counterregulatory response to the inhibition of intracellular glycolysis (Fig. 1I). As anticipated, we found that plasma GDF15 concentrations also increased after treatment with 2DG (Fig. 1J). Those data showed that GDF15 is an insulin-independent signal that is activated in the context of glucose shortage.

### GDF15 is produced by the kidney during hypoglycemia

Next, to confirm the source of GDF15 during hypoglycemia, we assessed the *Gdf15* mRNA expression in various tissues. The liver is a hub of glucose metabolism and is a source of GDF15 in a variety of situations including following metformin treatment, caloric excess^12^, amino acid deficiency^11^, and exercise^13^, as well as in non-alcoholic fatty liver disease^14, 15^ and lipodystrophy ^16^. However, in our mice, liver *Gdf15* mRNA expression was similar between mice treated with insulin and with phosphate-buffered saline (PBS). In contrast, *Gdf15* mRNA significantly increased in the kidney, but not in any other of the surveyed organs (Fig. 2A). In accordance with the transcript profile, GDF15 protein concentrations increased in the kidney but not in the liver (Fig. 2B). To localize the site of GDF15 production in the kidney, we utilized GDF15^GFP-Cre; R26^ tdTomato reporter mice. Compared to the PBS-treated mice, more cells in the medulla of the kidney in the insulin-treated mice showed the tdTomato signal (Fig. 2C). Next, to validate the source of GDF15, we employed fluorescence-activated cell sorting (FACS) to collect the tdTomato positive cells following treatment with insulin, and used bulk RNA sequencing to assess the transcriptome signature of GDF15-producing cells. The CIBERSORTx^17^ analysis revealed that GDF15 was primarily produced in the proximal tubule in hypoglycemic mice (Fig. 2D, Supplemental Fig. 2A-B). To confirm the segment of the renal proximal tubule in which GDF15 is produced during hypoglycemia, we assessed the colocalization of GDF15 with markers of segments of the nephron. Aquaporin-2 (AQP2) which is expressed at the collecting duct, did not co-localize with the tdTomato positive cells (Supplemental Fig. 2C). In addition, we did not observe co-localization of the tdTomato signal and SGLT2 or GLUT1 (Supplemental Fig. 2B), indicating that the early proximal tubule is not the source of GDF15. However, Megalin, a marker of the proximal tubule which is expressed at higher levels in the distal segments of the proximal tubule^18^, does co-localize with tdTomato staining (Fig. 2E). The expression pattern of the sodium-potassium ATPase in GDF15-expressing cells strongly suggests that GDF15 is produced in the S3 segment of the proximal tubule^19^ (Fig. 2F).

**Figure 2.**
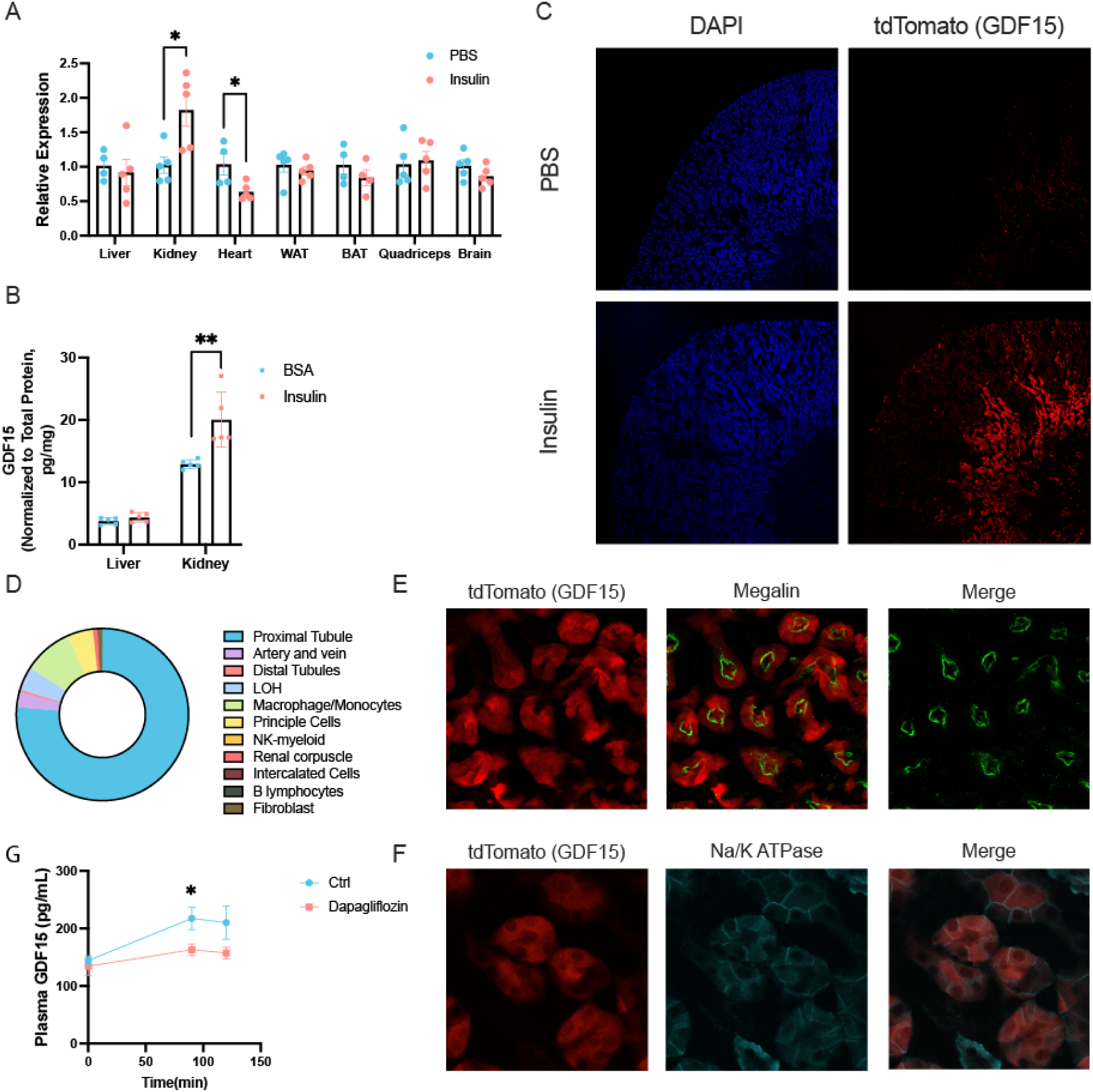
The S3 segment of the kidney produces GDF15 during hypoglycemia. (A) GDF15 mRNA expression in euglycemic (PBS-treated) or hypoglycemic (insulin-treated) mice. (B) GDF15 protein content in liver and kidney. (C) GDF15 tdTomato reporter mice demonstrate that GDF15 is produced in the kidney during hypoglycemia. (D) RNA sequencing data confirm that GDF15 is produced primarily in the renal proximal tubule during hypoglycemia in mice. (E) GDF15 is produced in the proximal tubule, as indicated by the colocalization of GDF15 tdTomato and megalin staining. (F) The S3 segment of the proximal tubule is the primary source of GDF15 during hypoglycemia, as indicated by the colocalization of sodium-potassium ATPase staining with GDF15 in tdTomato reporter mice. (G) Inhibiting glucose reabsorption in the proximal proximal tubule with dapagliflozin abrogates the secretion of GDF15 during an insulin tolerance test. In all panels, \**P*<0.05, \*\**P*<0.01 by the 2-tailed unpaired Student’s t-test.

Thus, we hypothesize that in a healthy mouse, most of the glucose in the renal filtrate is reabsorbed at the S1 and S2 segments of the proximal tubules, and the S3 segment reabsorbs the rest. However, during severe hypoglycemia, the S1 and S2 segments of the proximal tubule should reabsorb all of the glucose in the filtrate. The S3 segment then senses the absence of glucose and produces GDF15. To test this hypothesis, we treated the mice with the SGLT2 inhibitor dapagliflozin to inhibit the glucose reabsorption by S1 and, to a lesser extent S2 segments, presenting more glucose to the S3 segment. We found that dapagliflozin inhibits the production of GDF15 during an ITT (Fig. 2F).

### GDF15 defends against hypoglycemia by increasing gluconeogenesis

Having observed an increase in plasma GDF15 during both starvation and insulin-induced hypoglycemia in mice and rats, we aimed to determine whether the role of GDF15 in this setting is to serve as a hypoglycemia counterregulatory factor. Treatment with recombinant GDF15 (rGDF15) during an ITT modestly increased blood glucose concentrations (Fig. 3A). Blocking GDF15 prior to a hypoglycemic clamp increased the exogenous glucose required to maintain blood glucose concentrations between 40 and 60 mg/dL, whereas rGDF15 reduced the glucose infusion rate (Fig. 3C). The primary physiologic defense against hypoglycemia is an increase in endogenous glucose production (i.e. hepatic gluconeogenesis and glycogenolysis, and renal gluconeogenesis). We detected a reduction in endogenous glucose production in mice treated with anti-GDF15, and an increase following rGDF15 (Fig. 3D). After an overnight fast in rodents, glycogen is effectively depleted ^20, 21^ and gluconeogenesis is the sole contributor to endogenous glucose production. Utilizing stable isotope tracer methodology^22^, we found that most gluconeogenesis was fueled by phosphoenolpyruvate (PEP) in each group of mice. With a similar insulin concentration (Supplemental Fig. 3), rates of gluconeogenesis from PEP increased with rGDF15 and decreased with anti-GDF15 (Fig. 3E). In contrast, rGD15 did not alter hepatic glycogenolysis (Fig. 3F).

**Figure 3.**
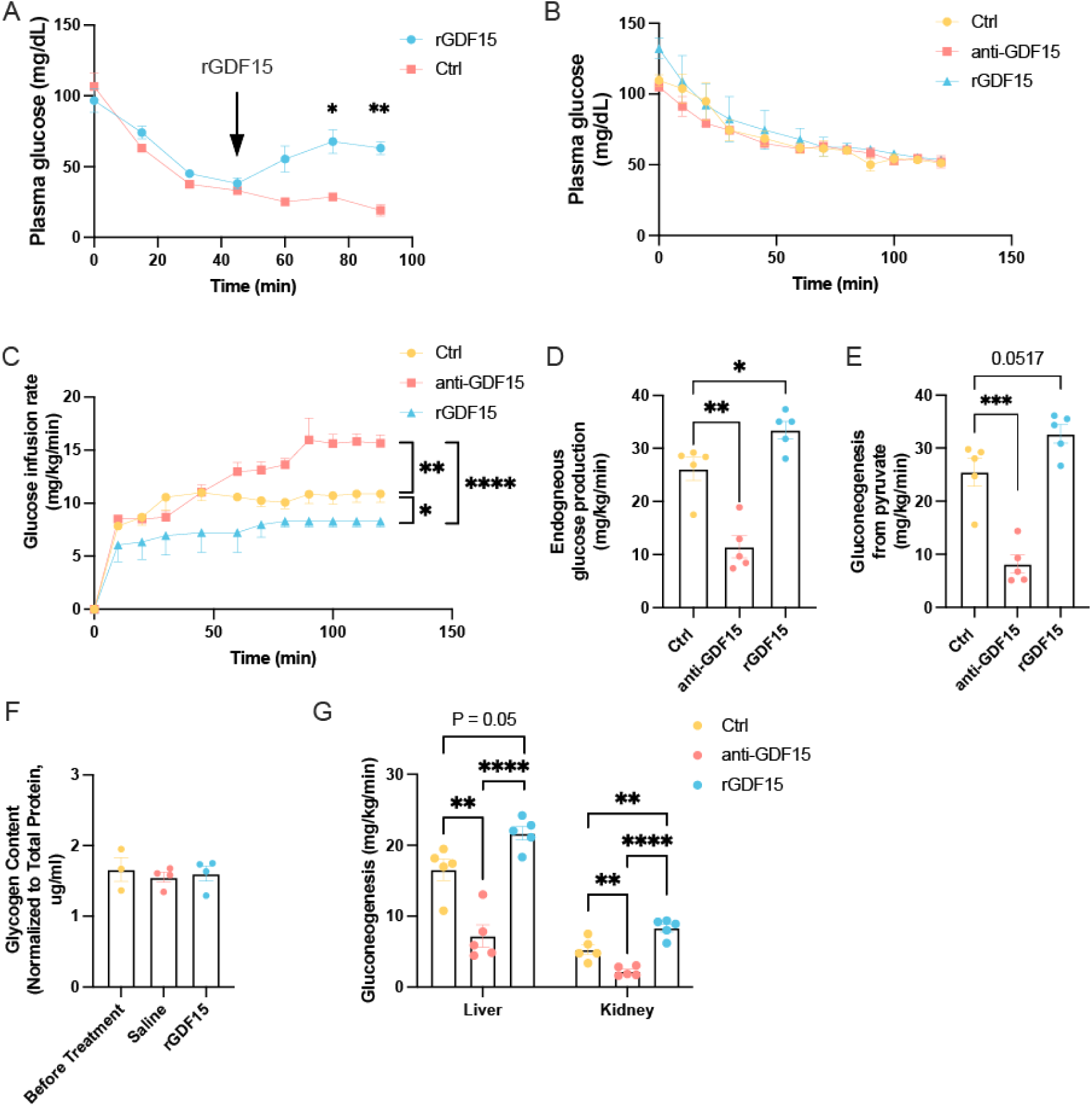
GDF15 defends against hypoglycemia by increasing gluconeogenesis from pyruvate. (A) Recombinant GDF15 administration increases plasma glucose concentrations during an insulin tolerance test. (B) Plasma glucose concentrations did not differ during a hypoglycemic clamp in mice treated with GDF15 neutralizing antibody or with recombinant GDF15, but (C) Recombinant GDF15 decreased, and anti-GDF15 antibody increased the glucose infusion rate during the hypoglycemic clamp. (D)-(E) Anti-GDF15 decreased, and recombinant GDF15 increased endogenous glucose production and gluconeogenesis from pyruvate. (F) GDF15 manipulations did not alter liver glycogen content. (G) GDF15 increased both hepatic and renal gluconeogenesis during a hypoglycemic clamp. In all panels, \**P*<0.05, \*\**P*<0.01, \*\*\**P*<0.001, \*\*\*\**P*<0.0001 by ANOVA with Tukey’s multiple comparisons test.

These data demonstrate that GDF15’s primary hypoglycemia counterregulatory mechanism is to stimulate gluconeogenesis. To determine the source(s) of these changes in gluconeogenesis, we employed Renal Gluconeogenesis Analytical Leads (REGAL)^23, 24^ to distinguish the contributions of liver and kidney to whole-body glucose production. GDF15 upregulated, and anti-GDF15 inhibited, gluconeogenesis from both liver and kidney (Fig. 3F). Because the liver produced the majority of glucose during hypoglycemia, regardless of the GDF15 manipulations, we elected to study the mechanism by which GDF15 promotes liver gluconeogenesis in the following experiments.

### GDF15 increases gluconeogenesis by increasing intrahepatic lipolysis

The primary mechanism of acute regulation of gluconeogenesis – on a scale of minutes, as is required for the immediate counterregulatory response to insulin-induced hypoglycemia – is allosteric. Acetyl-CoA, the endproduct of β-oxidation, is an allosteric activator of pyruvate carboxylase (PC)^25–28^, a rate-limiting gluconeogenic enzyme. Concentrations of acetyl-CoA may be increased by either increased white adipose tissue lipolysis^27^ and/or increased intrahepatic lipolysis^21^. Surprisingly, whole-body lipolysis as assessed by dilution of ^13^C palmitate tracer was not regulated by GDF15 manipulation (Fig. 4A). However, concentrations of both intrahepatic acetyl- and long-chain acyl-CoA were increased by GDF15 treatment and deceased by GDF15 blockade (Fig. 4B-C), which indicates that GDF15 is a regulator of intrahepatic lipolysis. Therefore, like glucagon^21^, we hypothesized that GDF15 can regulate gluconeogenesis by increasing intrahepatic lipolysis. To confirm if GDF15 stimulates gluconeogenesis via adipose triglyceride lipase (ATGL) activation, we generated liver-specific ATGL knockout (Atgl^f/f; Alb-CreER^) mice and treated the mice with rGDF15. The Cre-positive mice showed a significantly lower basal plasma glucose than did their Cre-negative littermates (Fig. 4D). During the hypoglycemic clamp, rGDF15 can only increase the endogenous glucose production in the WT mice, but not in ATGL-deficient mice (Fig. 4E).

**Figure 4.**
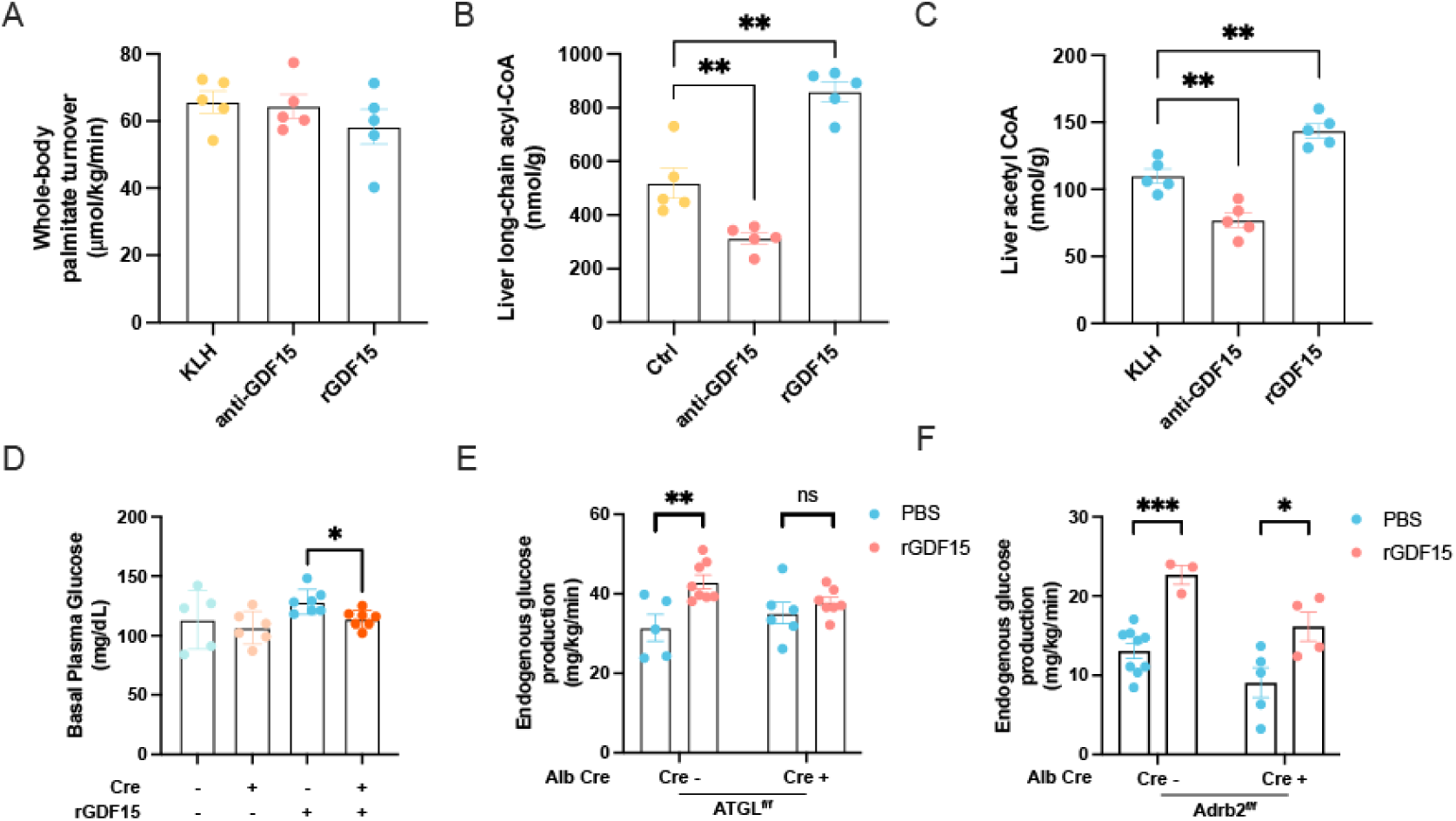
GDF15 promotes endogenous glucose production by increasing intrarenal lipolysis in an Adrb2-dependent manner. (A) Whole-body palmitate turnover did not differ with recombinant GDF-15 or anti-GDF15 treatment; however, both (B) Long-chain acyl- and (C) Acetyl-CoA concentrations in the liver were increased with recombinant GDF15 and reduced with anti-GDF15 treatment. (D) Mice deficient in intrahepatic lipolysis failed to induce gluconeogenesis in response to GDF15 treatment, as reflected by a reduced glucose infusion rate and (5) Reduced endogenous glucose production during a hypoglycemic clamp. (F) rGDF15 tends to induce liver and kidney epinephrine and norepinephrine. (H) Mice deficient in the liver Adrb2 showed poor induction of glucose production by rGDF15.

### GDF15 increases endogenous glucose production by the β-2 Adrenergic pathway

Β-adrenergic stimulation activates lipolysis in primary rat hepatocytes and human hepatoma cells^29^. The primary β-adrenergic receptors responsible for lipolysis and its consequent glucoregulatory effects are Adrb2 and Adrb3, but because Adrb3 is not found in liver^30^, we chose to focus on Adrb2. Therefore, we hypothesized that GDF15 increases intrahepatic lipolysis stimulating Adrb2 activity during hypoglycemia. To test this hypothesis, we performed hypoglycemic clamps in liver-specific Adrb2 knockout (Adrb2 ^f/f; Alb-CreERT2^) mice. Compared to their Cre-littermates, the Cre+ mice tend to have a lower endogenous glucose production (Fig. 4F), indicating an unsurprising defect in hypoglycemia counterregulation. To study whether the function of GDF15 is dependent on Adrb2 signaling, we treated the mice with rGDF15 prior to the hypoglycemic clamp. The exogenous rGDF15 can upregulate gluconeogenesis in the Cre-mice with a greater magnitude, as compared to Cre+ mice (Fig. 4F). These data show that GDF15 can regulate hepatic endogenous glucose production in an Adrb2-dependent manner.

### Mice with T1D exhibit impaired GDF15 production and GDF15 resistance

Next, we aimed to determine whether the newly identified role for GDF15 as a hypoglycemia counterregulatory hormone is relevant in type 1 diabetes (T1D). Most patients with tightly-controlled T1D exhibit not one but repeated episodes of hypoglycemia; therefore, we firstly studied if recurrent hypoglycemia *per se* affects the production of GDF15 upon subsequent hypoglycemia. We treated healthy mice with insulin daily for 3 days and assessed their GDF15 production during an ITT thereafter. The recurrent hypoglycemic mice showed similar glucose levels compared to the control group (Fig. 5A); however, they were less able to produce GDF15 during the ITT (Fig. 5B). In accordance with ITT, recurrent hypoglycemic mice showed lower GDF15 production during clamp (Fig. 5C). Similarly, the hypoglycemic clamp shows that the recurrent hypoglycemic mice have lower endogenous glucose production (Fig. 5C-D) and gluconeogenesis from PEP (Fig. 5F), reflecting an impaired hypoglycemia counterregulatory response.

**Figure 5.**
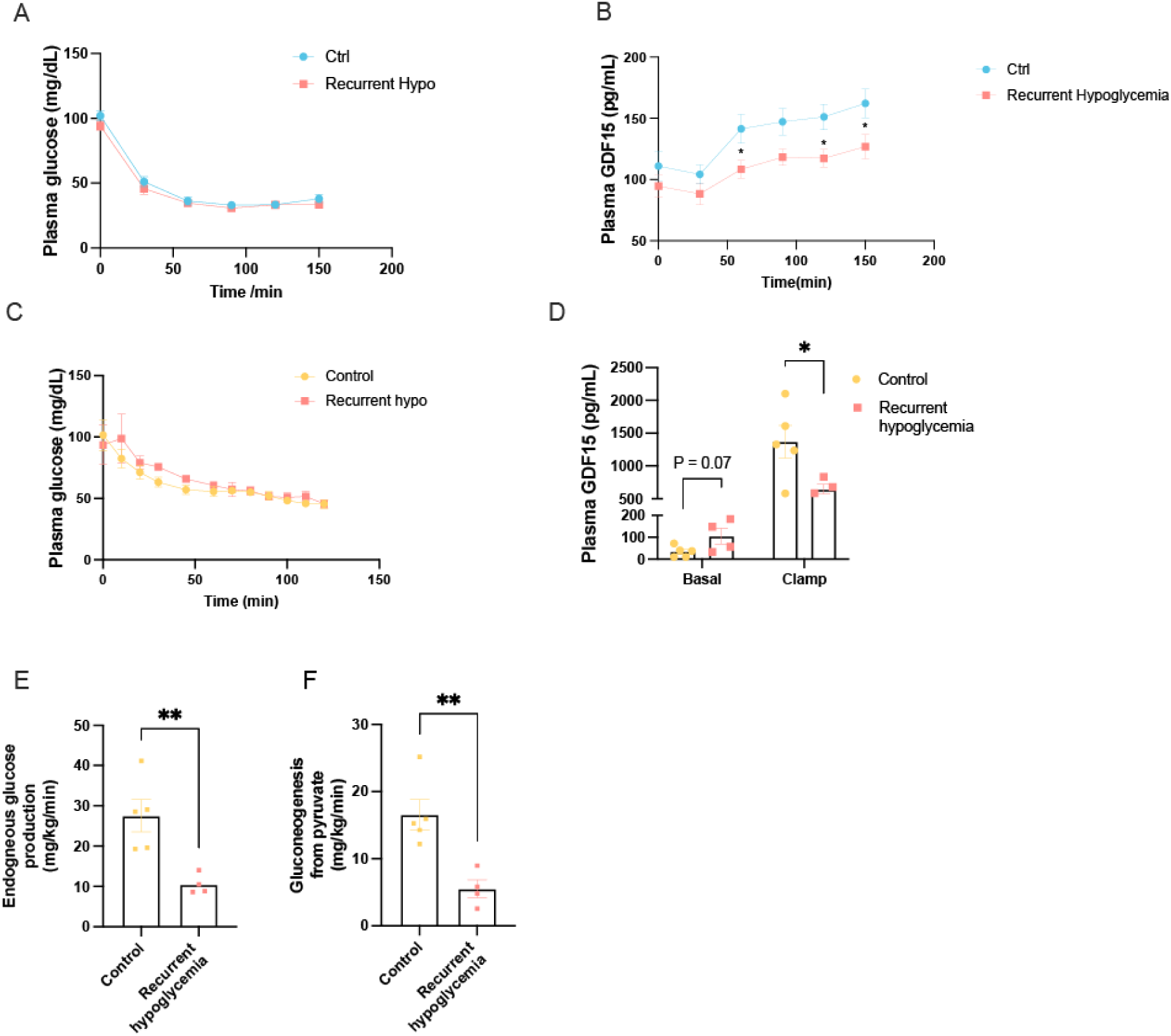
Recurrent hypoglycemia impairs GDF15 production in subsequent hypoglycemia. (A)-(B) Mice that experienced recurrent hypoglycemia for three days prior to the study exhibited impaired GDF15 secretion despite unchanged blood glucose concentrations during an insulin tolerance test. (C) Plasma GDF15 concentrations during a hypoglycemic clamp. (D)-(E) Endogenous glucose production and gluconeogenesis from pyruvate during the clamp. In all panels, \**P*<0.05, \*\**P*<0.01.

To study the impact of the pancreatic beta-cell loss on the GDF15 signal, we treated mice with streptozotocin to acutely induce T1D. The T1D mice showed a higher basal plasma GDF15 concentration and a more pronounced induction of GDF15 during the ITT (Fig. 6A-B). These data suggest that the T1D mice are resistant to the GDF15 signal. Consistent with this and unlike the healthy mice, T1D animals were resistant to exogenous rGDF15 and failed to increase blood glucose (Fig. 6C). In contrast, hypoglycemia in T1D mice can be rescued by the Adrb2 agonist clenbuterol (Fig. 6D), which indicates that GDF15 resistance occurs upstream of adrenergic outflow in mice with T1D.

**Figure 6.**
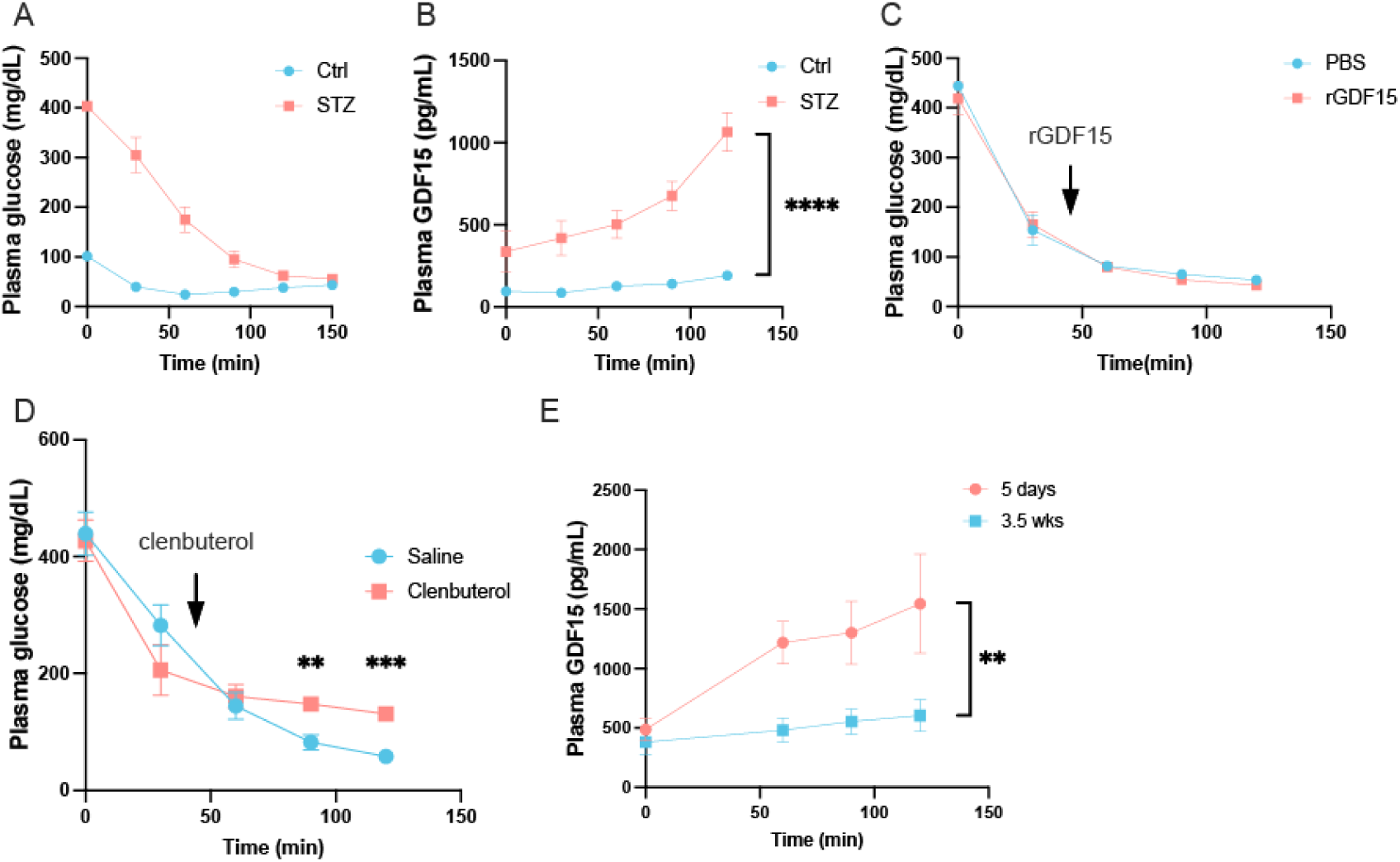
Mice with type 1 diabetes exhibit GDF15 resistance. (A) Streptozotocin-treated T1D mice are, as expected, hyperglycemic at baseline; however, when treated with high-dose insulin they achieve the same degree of hypoglycemia as healthy controls. (B) Plasma GDF15 is increased in T1D mice during the insulin tolerance test. (C) T1D mice fail to respond to recombinant GDF15 by increasing blood glucose during the ITT; however, (D) They do exhibit an increase in plasma glucose when treated with the Adrb2 antagonist clenbuterol, demonstrating that the defect is upstream of Adrb2. (E) GDF15 concentrations during an ITT in mice treated with STZ and either studied acutely, or maintained on subcutaneous insulin for 25 days.

Our T1D model differs from the clinical situation of most patients with T1D with regard to the duration of diabetes. Whereas mice, if untreated, must be used within one week of β-cell destruction with STZ, patients are maintained on insulin for years. To model the impact of established, insulin-treated diabetes on GDF15 production, we implanted insulin-releasing subcutaneous pellets in the T1D mice and compared the GDF15 produced during an ITT between untreated diabetic mice and mice treated with insulin pumps for 3.5 weeks. Insulin-treated diabetic mice produced significantly less GDF15 in terms of both absolute concentration and relative amplitude (Fig. 6E).

### Recurrent hypoglycemia causes impaired GDF15 production and glucose counter-regulation

Finally, we aimed to determine whether our results in mice would translate to humans. We performed hypoglycemic clamps in T1D patients and healthy controls who were matched for all relevant demographics and clinical characteristics, including age, sex, ethnicity, and body mass index (Table 1). As expected, patients with T1D exhibited lower fasting c-peptide and higher hemoglobin A1c. GDF15 was robustly induced during the hypoglycemic clamp in the control group, but not in T1D patients (Figure 7). These data confirm the translational relevance of the findings in this manuscript: that GDF15 is induced as a hypoglycemia counterregulatory factor in healthy individuals but not those with T1D.

**Figure 7.**
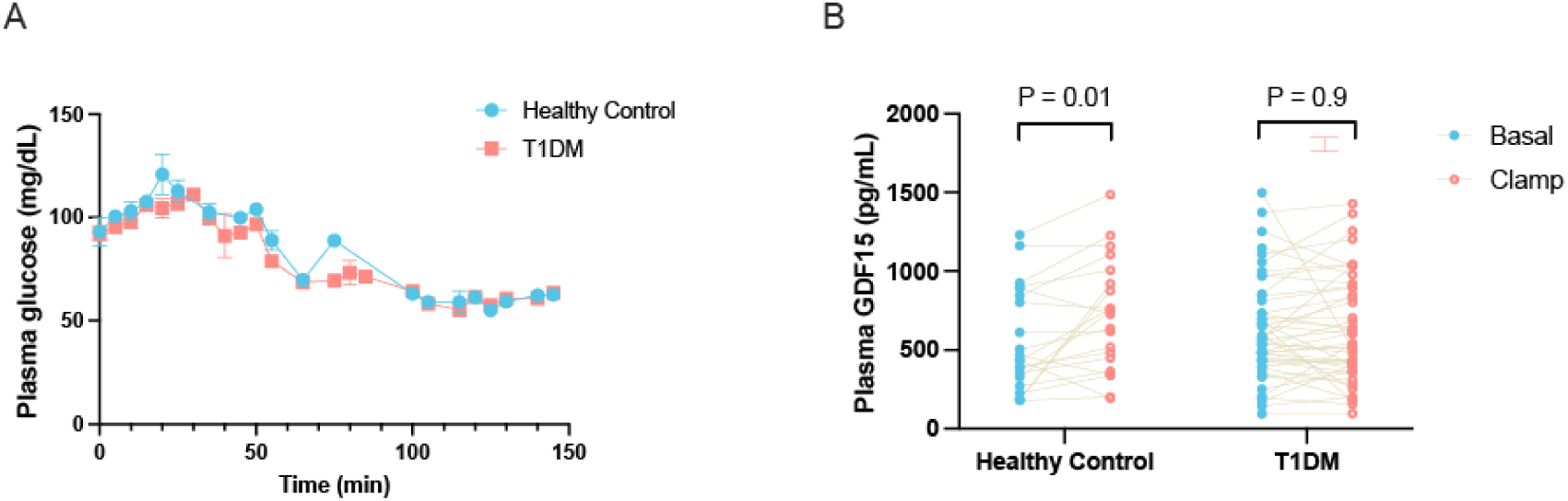
Patients with T1D exhibit impaired GDF15 secretion during a hypoglycemic clamp. (A) Plasma glucose concentrations throughout the clamp, which began at time zero). (B) Plasma GDF15 concentrations. Within-group comparisons were performed using the 2-tailed paired Student’s t-test. 21 healthy control volunteers and 47 subjects with T1D were studied.

**Table 1.** Clinical characteristics of participants in the human hypoglycemic clamp study. For continuous variables, the mean±S.E.M. is shown. \**P*<0.05, \*\**P*<0.01, \*\*\*\**P*<0.0001.

|  | Healthy Controls | Patients with T1D |
| --- | --- | --- |
| Number | 21 | 47 |
| Duration of diabetes (years) | N/A | 18.4±1.6 |
| Awareness (aware/unaware) | N/A | 28/19 |
| Gender (M/F) | 10/11 | 21/26 |
| Age (years) | 32±2 | 35±2 |
| Education (years) | 16.7±2.4 | 16.8±0.5 |
| BMI (kg/m<sup>2</sup>) | 24.3±2.9 | 25.8±0.7 |
| HbA1c (%) | 5.1±0.3 | 7.3±0.1**** |
| Basal c-peptide (ng/mL) | 1.71±1.00 | 0.21±0.03**** |
| Basal insulin (μU/mL) | 11.3±1.3 | 35.6±8.3 |
| Basal glucose (mg/dL) | 89±3 | 122±6** |
| Basal glucagon (pg/mL) | 49.6±8.5 | 32.5±3.9* |

## Discussion

GDF15 has garnered a great deal of attention in metabolism studies in recent years, focusing primarily on the chronic effects of GDF15. It has generally been considered a catabolic hormone, as shown by its role in increasing heat production and oxygen consumption ^31^. However, here we report an additional role for GDF15 as an anabolic hormone, increasing the glucose output from liver and kidney during hypoglycemia. We found that GDF15 can be produced by the S3 segment of the proximal tubule, consistent with other studies emphasizing the ability of the kidney ^32^ and, specifically, proximal tubules^33^ to produce GDF15 under very different conditions (acute kidney injury and Cockayne syndrome). However, to our knowledge, our data are the first to reveal GDF15’s role as a hypoglycemia counterregulatory hormone.

By employing the ITT and hypoglycemic clamp, we studied two translationally relevant scenarios of hypoglycemia. In both cases, GDF15 can effectively upregulate levels of plasma glucose, indicating its counterregulatory role in glucose control in a liver adrenergic signal and intrahepatic lipolysis-dependent manner. GDF15 was previously shown to stimulate the liver triglyceride output during acute sepsis^9^. Combined with our findings, GDF15 seems to have an effect to mobilize lipids in the liver by increasing both TG output and lipolysis.

We previously reported that glucagon can increase gluconeogenesis by increasing intrahepatic lipolysis^21^, and here we demonstrate that GDF15 can regulate gluconeogenesis in a similar way. The non-selective β adrenoceptor agonist clenbuterol can upregulate the rat hepatocyte lipolysis by activating ATGL via the cAMP/PKA pathway^29^. Thus, cAMP/PKA may be one of the signaling pathways that coordinate the signal from Adrb2 and glucagon receptor to orchestrate intrahepatic lipid metabolism.

Evolutionarily, fasting is one of the main challenges that animals must be prepared to withstand, so animals with redundant mechanisms to handle energy shortages have improved fitness. Glucagon and, to a lesser extent, epinephrine primarily stimulate glycogenolysis, so it is beneficial to have a complementary hormone(s) that primarily stimulate gluconeogenesis. In addition, our data demonstrate that it is the kidney that senses the glucose change, in response to the absence of any glucose passing through the S3 segment of the proximal tubule, and the kidney produces GDF15 in response. This novel mechanism serves as a “backup” means to respond to hypoglycemia when the counterregulatory actions of the pancreas and adrenal gland are insufficient.

In patients with type 1 diabetes (T1D), previous hypoglycemia can reduce the defense against subsequent hypoglycemia ^34–36^: even one episode of hypoglycemia can substantially impair the counterregulatory response ^35, 36^. However, the mechanism for this is incompletely understood. Both hypoglycemia-induced glucagon and epinephrine secretion were attenuated in patients with diabetes ^37–39^. The attenuated humoral response can contribute to the compromised defense against hypoglycemia in patients with diabetes. In this study, we showed that recurrent hypoglycemia *per se* in WT mice, and longstanding diabetes in humans, can cause an impaired GDF15 response to hypoglycemia. These data provide evidence for an additional causative link between previous hypoglycemia and impaired glucose regulation. In both patients with T1D and recurrently hypoglycemic mice, we find the GDF15 response to hypoglycemia is attenuated, which may translate into the abnormal glucose counterregulation. The defect in GDF15 production is possibly the result of pathological glucose variation or a chronic “hyper-GDF15” state. Studying how glucose counterregulation is achieved acutely and how it fails following repeated hypoglycemia can help us understand how metabolic homeostasis is maintained and will shed new light on potential therapies for acute hypoglycemia or impaired awareness of hypoglycemia.

It has been previously reported that the chronic GDF15 injection can cause anorexia and weight loss in healthy, obese, and tumor-bearing mice^40, 41^. GDF15 was also found to be critical in metformin-induced weight loss in a manner dependent on the transcription factors ATF4 and CHOP ^42^. This effect of GDF15 seems to be contradictory to our findings; however, it should be noted that at high concentrations, the classic hypoglycemia counterregulatory hormone glucagon can also cause nausea^43, 44^. Thus, the putative anorectic effect of GDF15 does not disqualify it from consideration as a hypoglycemia counterregulatory hormone. The evolutionary purpose of hypoglycemia-induced GDF15 is likely to boost gluconeogenesis during fasting, when glucocorticoid concentrations are high. The effect of glucocorticoids to promote hyperphagia likely trump the anorectic impact of GDF15 or glucagon. As a metabolic stress hormone, the dual function of GDF may be beneficial evolutionarily: the animal may intake pathogens and toxins from the environment, so when the overall energy balance is not life-threatening, GDF15 can decrease food intake to prevent intake of further pathogens or toxins, at the same time, GDF15 will increase the glucose output to provide enough substrate for maintenance of homeostasis.

Taken together, this study identifies a new anabolic role for GDF15 as a hypoglycemia counterregulatory hormone. In healthy humans and mice, GDF15 is secreted during hypoglycemia and promotes gluconeogenesis in part by stimulation of Adrb2-dependent intrahepatic lipolysis. These data deepen our understanding of GDF15 biology while identifying a new renokine, and suggest that the evolutionary function of this hormone may relate to its role as a hypoglycemia counterregulatory factor.

## Acknowledgments

We thank members of the Perry and Wang labs for insightful discussions. We are grateful for advice on kidney physiology from members of the Caplan lab and Dr. Tong Wang. Kind assistance in the human hypoglycemic clamps was provided by Dr. Elizabeth Sanchez Rangel, Mari-Lynet Knight, and Jelani Deajon Jackson. Recombinant GDF15 and anti-GDF15 antibody were from NGM Biopharmaceuticals and as previously described^9, 10^ This study was funded by grants from the Juvenile Diabetes Research Foundation (R.J.P. and A.W.) and the U.S. Public Health Service (R37CA258261A1 [R.J.P.], R01AI162645 (A.W.), R01AR080104 (A.W.), RC2DK120534 (M.J.C.), R01DK072612 (M.J.C.), and R01DK020495 [J.J.H.]). Human samples were analyzed by the Yale Diabetes Research Center Clinical Metabolism core (P30 DK045735).

## Author Contributions

The study was conceived and designed by Z.L. and R.J.P. Experiments were performed and data analyzed by Z.L., X.Z., Q.X., C.Y., Q.Z., S.S., B.G.C.L., X.L., K.I.-W., A.R.N., M.J.C., R.B.C., R.B.A., J.J.H., A.W., and R.J.P. Mouse surgeries were performed by W.Z., and knockout mice generated and supplied by C.Z. and A.W.

## Star Methods

### RESOURCE AVAILABILITY

Further information and requests for resources and reagents should be directed to and will be fulfilled by the lead contact, Rachel Perry. This study did not generate new unique reagents. All data reported in this paper are shown in the dot plots and will be shared by the lead contact upon request. Any additional information required to reanalyze the data reported in this paper is also available from the lead contact upon request.

### EXPERIMENTAL MODEL AND SUBJECT DETAILS

#### Mice and Rats

The Yale Institutional Animal Care and Use Committee approved all studies prior to experimentation. Wild-type C57bl/6J mice were purchased from Jackson Laboratories (stock number 000664). Liver-specific ATGL knockout mice were generated by crossing Atgl^f/f^ mice (Jackson Labs #024278) with Alb-Cre animals (Jackson Labs #003574). Liver-specific Adrb2 knockout mice were generated by crossing Adrb2^f/f^ (donated by Gerard Karsenty from Columbia University) with Alb-Cre animals (donated by Nikhil Joshi from Yale University). GDF15^GFP-Cre; R26^ were donated by Andrew McMahon from the University of Southern California in which the promoter of GDF15 can induce the expression of Cre recombinase to remove the stop codon before the tdTomato gene. Genotyping was performed by PCR, using primers from IDT with sequences shown in Table 3. Wild-type Sprague Dawley rats were purchased from Charles River (strain code 400)

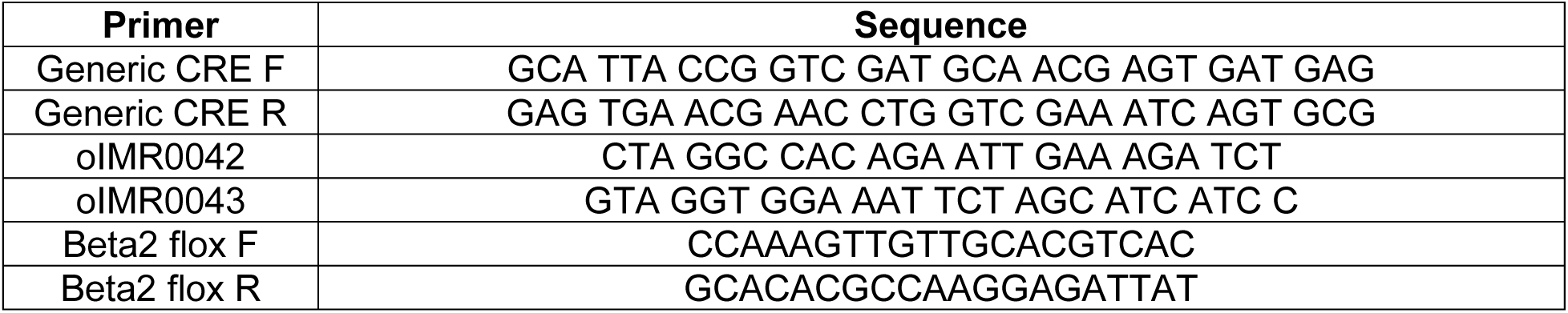

Male mice were studied between 7 and 12 weeks of age (body weight 19-25 grams), and rats at 10 weeks of age (body weight 280-320 grams). Mice were group housed prior to one week before the terminal study, when they underwent surgery (as detailed below) and were subsequently singly housed. In all pharmacologic studies, vehicle controls (as described in the figures and figure legends) were used, studied simultaneously with experimental animals. Specific measures to randomize animals to treatment were not undertaken; however, all studies were performed in littermate controls, with mice from each home cage randomized to each intervention group. In studies of genetically modified mice, wild-type littermates served as controls. Sample sizes for *in vivo* studies were determined to provide 95% power to detect an expected difference in endogenous glucose production (the primary outcome measure) of 30% with 30% standard deviations. Animals were excluded from further analysis if they did not immediately respond to intravenous pentobarbital, but otherwise no animals or data were excluded from analysis. All analyses were performed by investigators who were blinded to group allocation during the data analysis. It was not practically possible to blind investigators to group allocation during the experiments (e.g. hypoglycemic clamps or insulin tolerance tests).

#### Human Subjects

The human hypoglycemic clamp study was approved by the Yale Institutional Review Board. Adults over age 18 of both sexes (as shown in Table 1) were included. Informed consent was obtained prior to any intervention.

### METHOD DETAILS

#### Rodent Clamps

Rodents underwent surgery under isoflurane anesthesia to place silastic catheters in the right jugular vein (mice), and in the right jugular vein and left carotid artery (rats). Catheters were filled with saline containing heparin (10 U/mL), and were tunneled to the back of the head and tied off with a suture so that they were inaccessible to the animal. After surgery, mice were treated with carprofen (5 mg/kg) daily for three days. On day 7 after surgery, following an overnight (14 hour) fast, the catheters were opened by saline flush. Catheters were connected and animals were allowed to acclimate for two hours with free movement around their cage (rats) or in a plastic restrainer, gently tethered by taping the tail to the restrainer (mice).

During the hypoglycemic clamps, Regular insulin was infused continuously (4 mU/kg/min in mice, 2 mU/kg/min into the arterial catheter in rats). Artificial plasma^45^ (115 mM NaCl, 5.9 mM KCl, 1.2 mM MgCl_2_, 1.2 mM HaH_2_PO_4_, 1.2 mM Na_2_SO_4_, 2.5 mM CaCl_2_, 25 mM HaHCO_3_, 4% BSA, filtered through 0.22 um filtered and adjust to pH = 7.45) was used to dilute the tracers. In rats, serum from the studied animals was used to bind the insulin. Blood was collected by tail massage (mice) or through the venous catheter (rats) every 10-15 min, and 20% dextrose was infused at a variable rate to maintain euglycemia (110-120 mg/dL) or hypoglycemia (∼70 mg/dL in rats, 40∼60 mg/dL in mice). Concurrently, mice were infused with [3-^13^C] lactate (40 umol/kg/min) and [^2^H_7_] glucose (0.4 mg/kg/min). After 120 min of the clamp, animals were euthanized with intravenous Euthasol, and livers and kidneys freeze-clamped using tongs pre-chilled in liquid nitrogen. Samples were stored at -80°C for further analysis.

#### Other metabolic interventions in rodents

For the insulin tolerance tests, overnight fasted mice were injected IP with insulin (1 U/kg in healthy mice, 4 U/kg in T1D mice) Humulin (NDC 0002-8215-01) was diluted by 5% fatty-acid-free BSA in saline. Blood was collected by tail massage every 15 min for 90-150 minutes. In the fasting studies, mice were singly housed with food removed at 4:00 pm, and blood was collected every 24 hours. To generate type 1 diabetes, mice were injected with 180 mg/kg streptozotocin dissolved in the Citrate solution (pH = 4.5)^46^. The streptozotocin solution was freshly made for each experiment and used within 20 min after dissolving.

#### LC/MS Samples Preparation

30 ul 10% Methanol in Acetonitrile (w/w) buffer was added to 10 ul plasma to precipitate the protein. The mixture was centrifuged for 15 min at 10000 RPM. Then, the supernatant liquid was transferred to the filter{I will check} and centrifuged for 20 min at 10000 RPM. The flow-through was analyzed by Liquid Chromatography-Mass Spectrometry.

#### Flux analysis

Plasma ^13^C glucose enrichment was determined by gas chromatography/mass spectrometry^20^, and endogenous glucose turnover was calculated using the equation

Endogenous glucose production = [(Tracer ^13^C APE/Plasma ^13^C APE)-1]*Infusion rate where APE denotes the measured atom percent enrichment. We utilized our previously published method^22^ to measure gluconeogenesis from PEP in both liver and kidney, and applied our recently reported method to determinine hepatic versus renal gluconeogenesis^47^.

#### Pharmacologic interventions

In the hypoglycemic clamp, rGDF15 was administrated subcutaneously 1 hour before the start of the clamp. Anti-GDF15 antibody was injected intraperitoneally 15 hours before the experiment. 1 mg/kg Dapagliflozin was administrated by oral gavage 2 hours before the experiment. 2-Deoxy-d-glucose was injected intraperitoneally with a dose of 1g/kg.

#### Biochemical analysis

Blood glucose concentrations were measured with the iPet PRO handheld glucometer. Plasma GDF15 concentrations in mice and rats were measured by ELISA (R&D Systems). Glycogen concentration was measured using the phenol-sulfuric acid method^48^. Long-chain acyl^49^- and acetyl-CoA concentrations^27^ were measured by liquid chromatography-mass spectrometry/mass spectrometry.

#### Imaging studies

GDF15^GFP-Cre; R26^ tdTomato reporter mice were injected with 12.5 mg/kg tamoxifen at 5 pm 1 day prior to the experiment day and the second dose of tamoxifen was given at 10 am of the experiment day. ITT was performed with those mice at 2 pm and food was provided at 5 pm. 48 hours later, the liver and kidney were collected and saved in 4% PFA at 4 ℃ for 18 to 24 hours. Fixed samples were transferred into 20% sucrose (PBS) solution until sinking to the bottom. Then, samples were frozen in O.C.T. and cryo-sectioned into 10 um slices. Nephron markers were stained and imaged with Leica SP8 confocal microscope.

#### RNA sequencing

Mice were kept in cages for 2 days after ITT with *ad lib* access to regular chow. Then mice were sacrificed and the kidney was collected. The kidney was cut into small pieces and incubated in the digestion buffer containing 25 ug/ml Liberase TM and 50 µg/ml DNase in RPMI media. Samples were kept at 37℃ for 45 min and passed through a 100-um cell strainer. Single cells were isolated by FACS according to the tdTomato signaling and collected into the RLT buffer containing BME. Total mRNA was extracted by Qiagen RNAeasy kit and sequenced by BGI. Reads were aligned to mm39 and analyzed with CIBERSORTx^17^ in which single-cell sequencing data from Kidney Cell Explorer^18^ was used as the reference.

#### Human clamps

Participants were recruited from the greater New Haven area, some as part of a prior study^50^. Inclusion criteria included age 18-50, non-smoking, BMI ≥18.5, and no alcohol or drug use within 72 hours of the study, and exclusion criteria included illicit drug or recent steroid use, active infection, malignancy, abnormal thyroid function, cerebrovascular or cardiovascular disease, weight change in the last 3 months, and pregnancy or breastfeeding. Patients with type 1 diabetes with no history of neuropathy or proliferative retinopathy were recruited pursuant to the above inclusion/exclusion criteria. Prior to the study, all participants completed a screening visit in which they provided a brief medical history, and a venous blood sample was drawn to measure hematocrit, creatinine, hemoglobin A1c, and C-peptide. On the day of the study, following an overnight fast, antecubital catheters were placed bilaterally. One was used to deliver insulin (2 mU/kg/min) and a variable infusion rate of dextrose, and the other to draw blood. The basal blood draw was performed at time zero of the clamp and counterregulatory hormone concentrations at time zero were compared to those measured in the final clamp sample. Plasma GDF15 concentrations were measured by ELISA (R&D Systems), and insulin, c-peptide, glucagon, epinephrine, norepinephrine, and cortisol by the Yale Diabetes Research Center Clinical Metabolism core.

### QUANTIFICATION AND STATISTICAL ANALYSIS

The 2-tailed Student’s t-test (paired or unpaired) was used to compare two groups, and two-way ANOVA with Tukey’s multiple comparisons test to compare three or more groups. Statistical analyses were performed using GraphPad Prism version 9.

### KEY RESOURCES TABLE

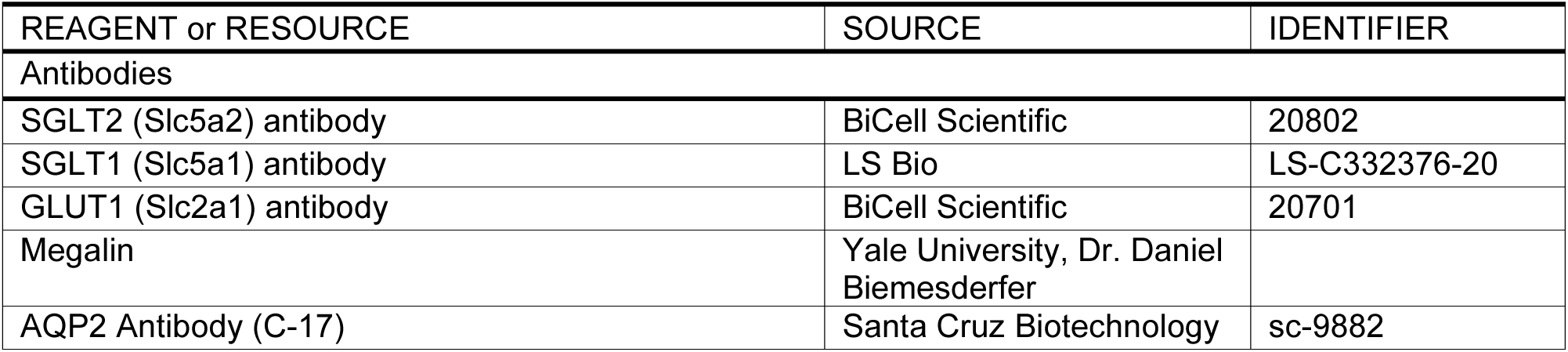
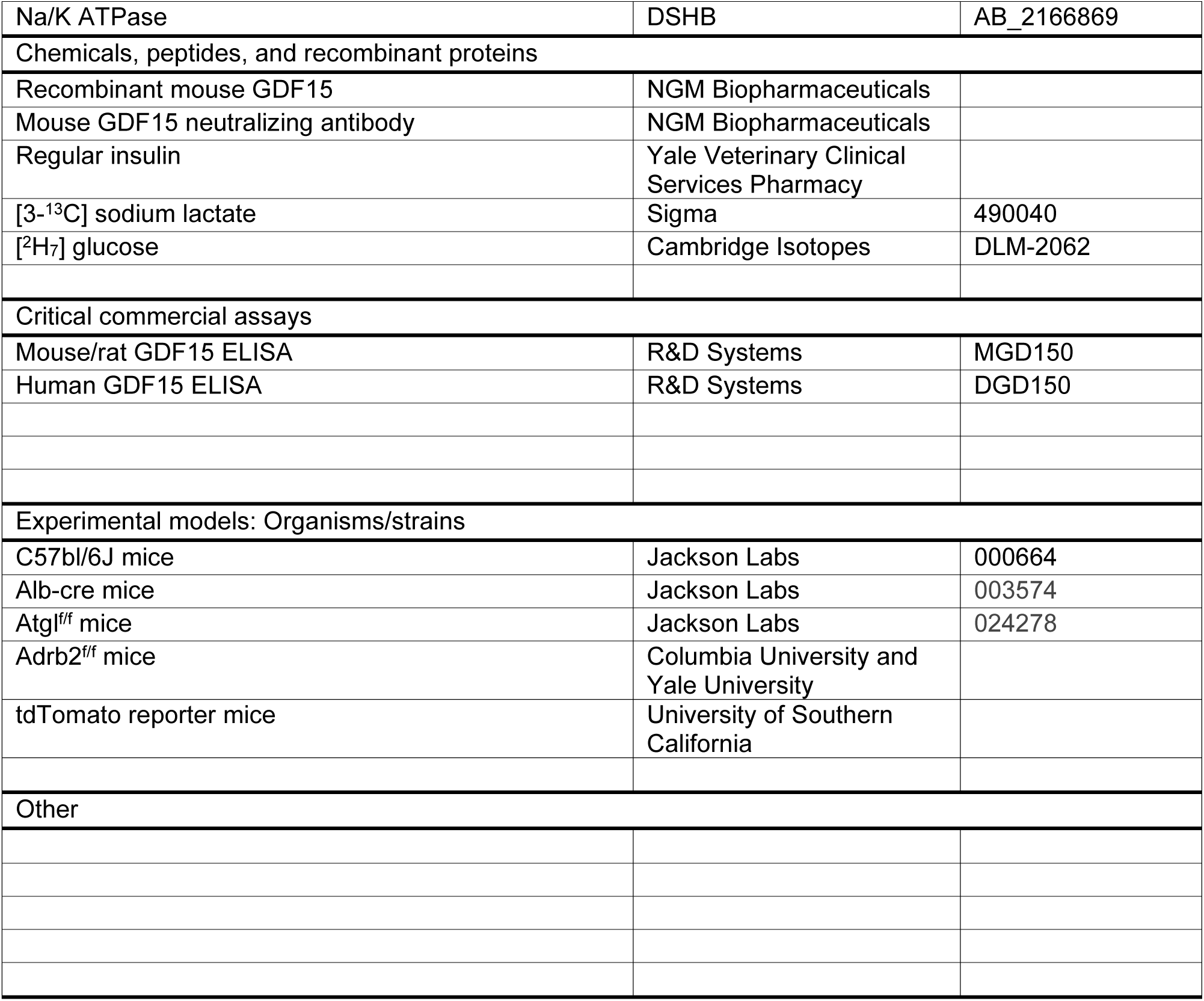

## Supplemental Data

**Supplemental Figure 1.**
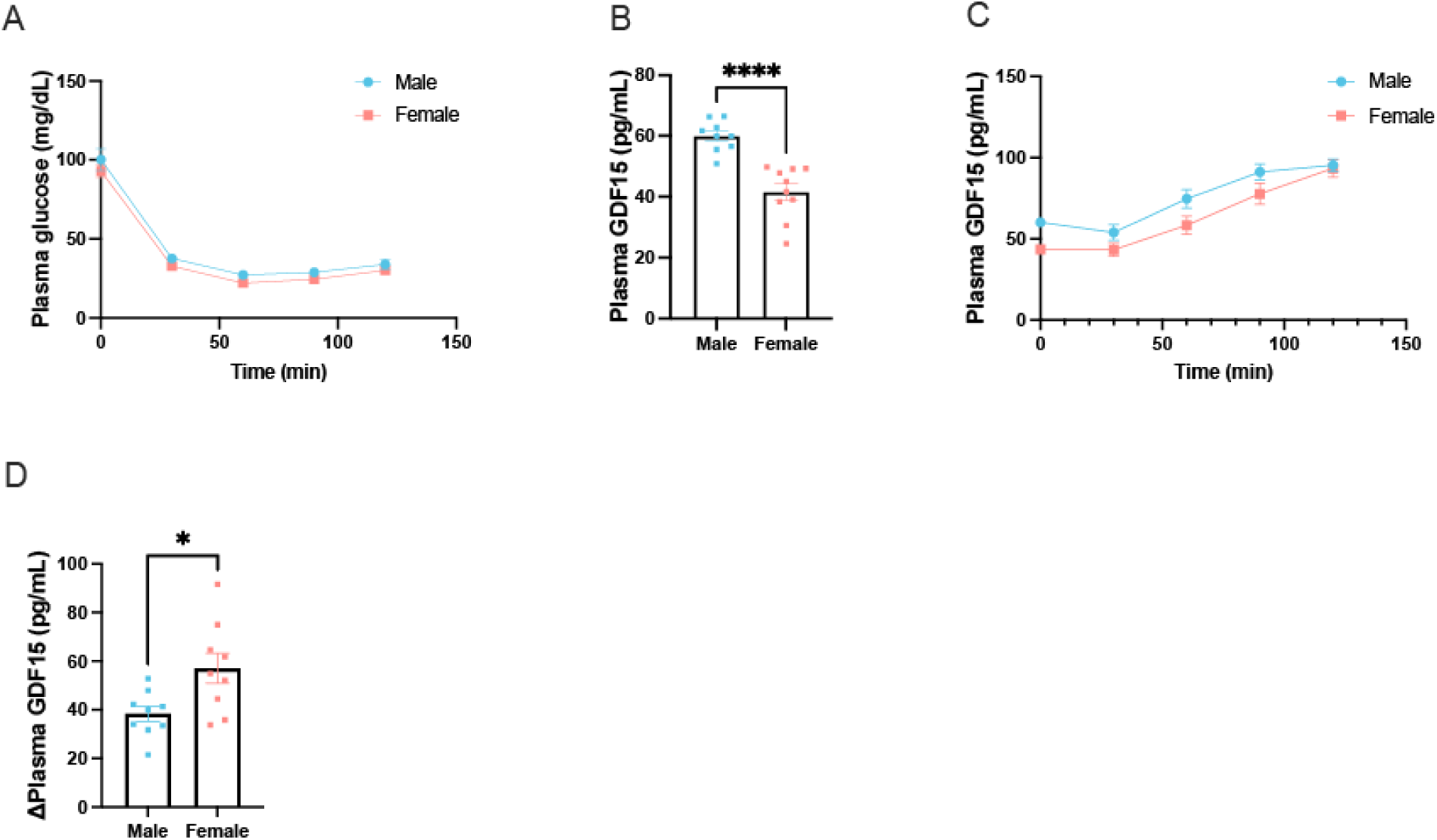
Sex differences in GDF15 induction in response to hypoglycemia. (A) Blood glucose during an ITT in male and female mice. (B) Basal plasma GDF15 and (C) GDF15 throughout the ITT. (D) Change in GDF15 from time zero to 120 min of the ITT. \**P*<0.05, \*\*\*\**P*<0.0001.

**Supplemental Figure 2.**
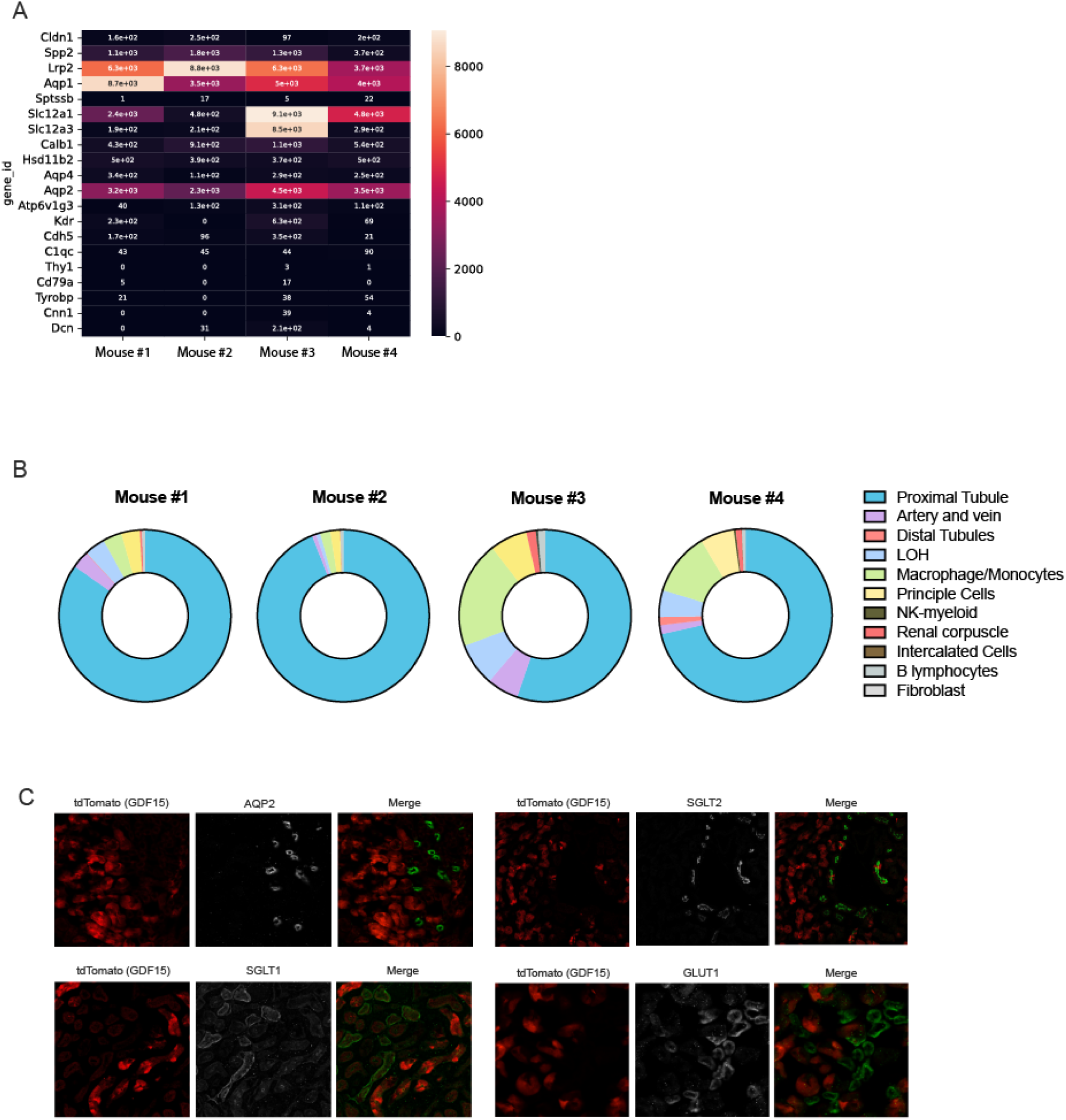
During hypoglycemia, GDF15 production is primarily induced in the renal proximal tubule. (A) Bulk RNA sequencing of the GDF15-expressing cells shows the signature of proximal tubules, confirmed by CIBERSORTx analysis. (B) CIBERSORTx analysis of the bulk RNA sequencing data from each individual mouse. (C) The proximal tubule identity was confirmed by the staining pattern of AQP2, SGLT1, SGLT2, and GLUT1.

**Supplemental Figure 3.**
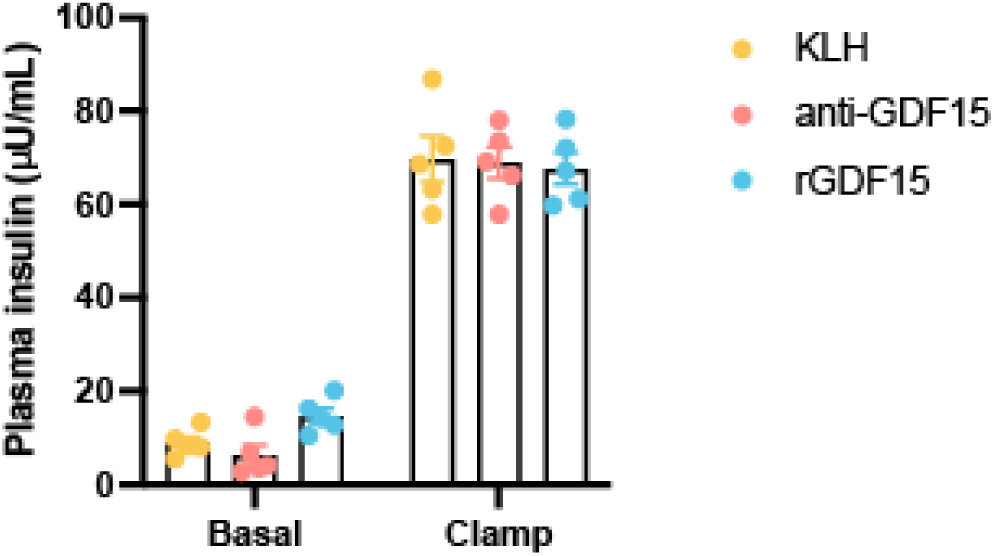
**Plasma insulin concentrations did not differ** in the basal or clamp settings between control (KLH-treated), anti-GDF15-treated, or recombinant GDF15-treated mice.

